# The Rpd3-complex regulates expression of multiple cell surface recycling factors in yeast

**DOI:** 10.1101/2021.10.22.465438

**Authors:** Konstantina Amoiradaki, Kate R Bunting, Katherine M Paine, Josephine E. Ayre, Karen Hogg, Kamilla ME Laidlaw, Chris MacDonald

**Affiliations:** York Biomedical Research Institute and Department of Biology, University of York, York, UK; Imaging and Cytometry Laboratory, Bioscience Technology Facility, Department of Biology, University of York, UK

## Abstract

Intracellular trafficking pathways control residency and bioactivity of integral membrane proteins at the cell surface. Upon internalisation, surface cargo proteins can be delivered back to the plasma membrane via endosomal recycling pathways. Recycling is thought to be controlled at the metabolic and transcriptional level, but such mechanisms are not fully understood. In yeast, recycling of surface proteins can be triggered by cargo deubiquitination and a series of molecular factors have been implicated in this trafficking. In this study, we follow up on the observation that many subunits of the Rpd3 lysine deacetylase complex are required for recycling. We validate ten Rpd3-complex subunits in recycling using two distinct assays and developed tools to quantify both. Fluorescently labelled Rpd3 localises to the nucleus and complements recycling defects, which we hypothesised were mediated by modulated expression of Rpd3 target gene(s). Bioinformatics implicated 32 candidates that function downstream of Rpd3, which were over-expressed and assessed for capacity to suppress recycling defects of *rpd3Δ* cells. This effort yielded 3 hits: Sit4, Dit1 and Ldb7, which were validated with a lipid dye recycling assay. Additionally, the essential phosphatidylinositol-4-kinase Pik1 was shown to have a role in recycling. We propose recycling is governed by Rpd3 at the transcriptional level via multiple downstream target genes.

## INTRODUCTION

Most integral membrane proteins expressed in eukaryotic cells are inserted into the endoplasmic reticulum (ER) via different mechanisms [1]. Many of these perform diverse roles at the plasma membrane (PM), such as ion channels, nutrient transporters, and different classes of receptors [2,3], and are actively transported from the ER through the secretory pathway to the cell surface [4]. Mechanisms of surface protein regulation have been elucidated using the budding yeast *Saccharomyces cerevisiae*, where hundreds of proteins are organised in distinct spatial arrangements [5]. The lateral movement of proteins between regions of the PM correlates with their biological activity [6]. For example, inactive nutrient transporters localised to eisosome subdomains adopt an active conformation for nutrient uptake upon migration to other regions of the PM in response to substrate [7,8]. This altered PM localisation of active transporters supports their internalisation and endocytosis [9], a process which is controlled metabolically, with stress conditions altering eisosomal capacity to harbour nutrient transporters [10,11].

Surface proteins are internalised to the endosomal system, a network of intracellular compartments that organise and traffic protein and lipid material to other intracellular destinations [12]. Surface membrane proteins destined for degradation are retained in endosomes, which undergo a maturation process to definable late endosomes called multivesicular bodies (MVBs), which interface with lysosomes to drive cargo degradation [13]. The ubiquitination of membrane proteins serves as a conserved signal for trafficking through the degradative MVB pathway [14]. However, studies in various systems have shown that cargo deubiquitination cancels the degradation signal and triggers surface recycling [15]. Surface cargo recycling back to the PM in mammalian cells occurs either directly or via distinct compartments [16]. In yeast, surface recycling of cargoes from Vps4-endosomes is triggered by cargo deubiquitination [17].

Fluorescently labelled lysosomal cargoes fused to the catalytic domain of a deubiquitinating enzyme (DUb) are rediverted back to the PM and serve as reporters for this deubiquitination-induced recycling pathway [18]. A GFP-tagged DUb-fusion of the G-protein coupled receptor (GPCR) Ste3, which localises exclusively to the surface in wild-type cells, was used to screen for potential recycling factors that mis-localise the reporter [19]. This assay was calibrated with recycling mutants lacking *RCY1*, which have defective trafficking of recycled material, including the yeast synaptobrevin Snc1, the GPCR Ste2 and lipids labelled with the amphiphilic styryl dye FM4-64 [20,21]. Characterisation of this yeast recycling pathway has shown requirement for the Rag GTPases [19], the endosomal sorting complexes required for transport (ESCRT)-related protein Ist1 [17], and the phosphatidylinositol 3-kinase activity effector Gpa1 [22]. However, whether the pathway is regulated at a transcriptional level, and what the downstream molecular players might be, is unknown.

Acetylation of protein substrates is a common co- or post-translational modification whereby an acetyl group is covalently attached to proteins at their N-terminus or lysine residues [23]. Protein acetylation can alter the behaviour, biological activity, and stability of modified protein substrates, and has been implicated in a range of human diseases [24]. Acetylation of lysine residues, performed by lysine acetyltransferase (KAT) enzymes, is a reversible process that is antagonised by various lysine deacetylases (KDAC) enzymes. These enzymes were initially shown to modify histones [25] by regulating chromatin condensation and transcriptional activity [26]. Although, there are many additional functional consequences of protein acetylation, in both eukaryote and prokaryote systems [27]. KDACs (or histone DACs, HDACs) in human cells can be classified into two groups, the seven members of the NAD^+^-dependent sirtuin family [28] and the eleven members of the ‘classical’ Rpd3/Hda1 family [29].

Rpd3 is a yeast KDAC that is highly conserved throughout evolution [30] and has predominantly been linked with transcriptional regulation [31–33]. A large series of genetic, biochemical, and proteomic efforts have robustly characterised Rpd3 interactions (~100 physical and ~1000 genetic) and shown Rpd3 exists in two main complexes, termed Large (Rpd3L) and Small (Rpd3S), which have functionally distinct actions [30]. Almost all Rpd3 subunits localise to the nucleus with a small cytosolic population [34–37], where transcriptional control occurs via histone modification. Multiple members of the Rpd3-complex have been implicated in various biological processes, such as chromatin stability [38], DNA damage [39], drug sensitivity [40,41], and physiological stress responses [42,43]. Rpd3 was also shown to be required for efficient recycling of cargoes from endosomes back to the PM [19], as discussed above. In this study, we examined ten Rpd3-complex members that were previously implicated in yeast recycling and identify the downstream molecular factors that mediate recycling.

## RESULTS

### The Rpd3 complex is required for efficient surface recycling

Endosomal recycling can be tracked using Ste3-GFP-DUb [18], which recycles efficiently in wild-type cells but is retained in intracellular endosomes in recycling mutants such as *rcy1Δ* cells (**Figure 1A**), demonstrated by 3D confocal projections of the recycling reporter in wild-type cells and recycling defective *rcy1Δ* mutants (**Movie S1**). Plasmid expression of Ste3-GFP-DUb in 4,985 haploid deletion mutants revealed 89 validated mutants that are defective in cell surface recycling. Amongst this list of potential recycling factors were *rpd3Δ* cells, which lacks a histone modifying regulator of gene expression [33], alongside 9 other mutants of the Rpd3-complex [44] (**Figure 1B**). To confirm these results, we stably integrated Ste3-GFP-DUb into wild-type and all 10 previous identified mutants of the Rpd3 complex, revealing all mutants had some degree of recycling inefficiency, with intracellular accumulation of reporter similar to that observed in *rcy1Δ* cells (**Figure 1C**). To quantify differences between different null strains lacking component of the Rpd3-complex, we optimised the segmentation of cells using phase contrast (PC) and digital interference contrast (DIC). Although PC introduced an obvious border to define cells for segmentation in brightfield micrographs, we noted a significantly poorer fluorescence signal (**Figure 2A**). We therefore refined parameters to define cells from DIC images and used this protocol to estimate the background autofluorescence in the green channel capturing Ste3-GFP-DUb expressing yeast cells. For this, cells expressing Ste3-GFP-DUb were mixed with a separate culture of cells expressing Gpa2-mCherry and imaged (**Figure 2B**). Gpa2 is a Gα-subunit that regulates cAMP production that exclusively localises to the periphery via lipid modifications [22,45]. Segmentation of cells expressing Gpa2-mCherry allowed cellular autofluorescence to be measured in optical conditions optimised for Ste3-GFP-DUb acquisitions (**Figure 2C**). Having optimised segmentation and normalisation parameters, we then used a morphological erosion function [46] to measure the levels of cell surface signal as a percentage of total fluorescence. As expected, Ste3-GFP-DUb and Gpa2-mCherry, which both primarily localise to the PM at steady state, had similarly high levels of PM localisation (**Figure 2D**). Applying this analysis to all mutants of the Rpd3-complex expressing Ste3-GFP-DUb revealed every mutant had defective recycling compared to wild-type cells (**Figure 2E**). We found the most defective mutant was *hos2Δ*, which was as defective as the prototypical recycling mutant strain *rcy1Δ*. The least defective mutant was *ume1Δ*, which still exhibited significantly impaired recycling compared to wild-type cells.

**Figure 1:**
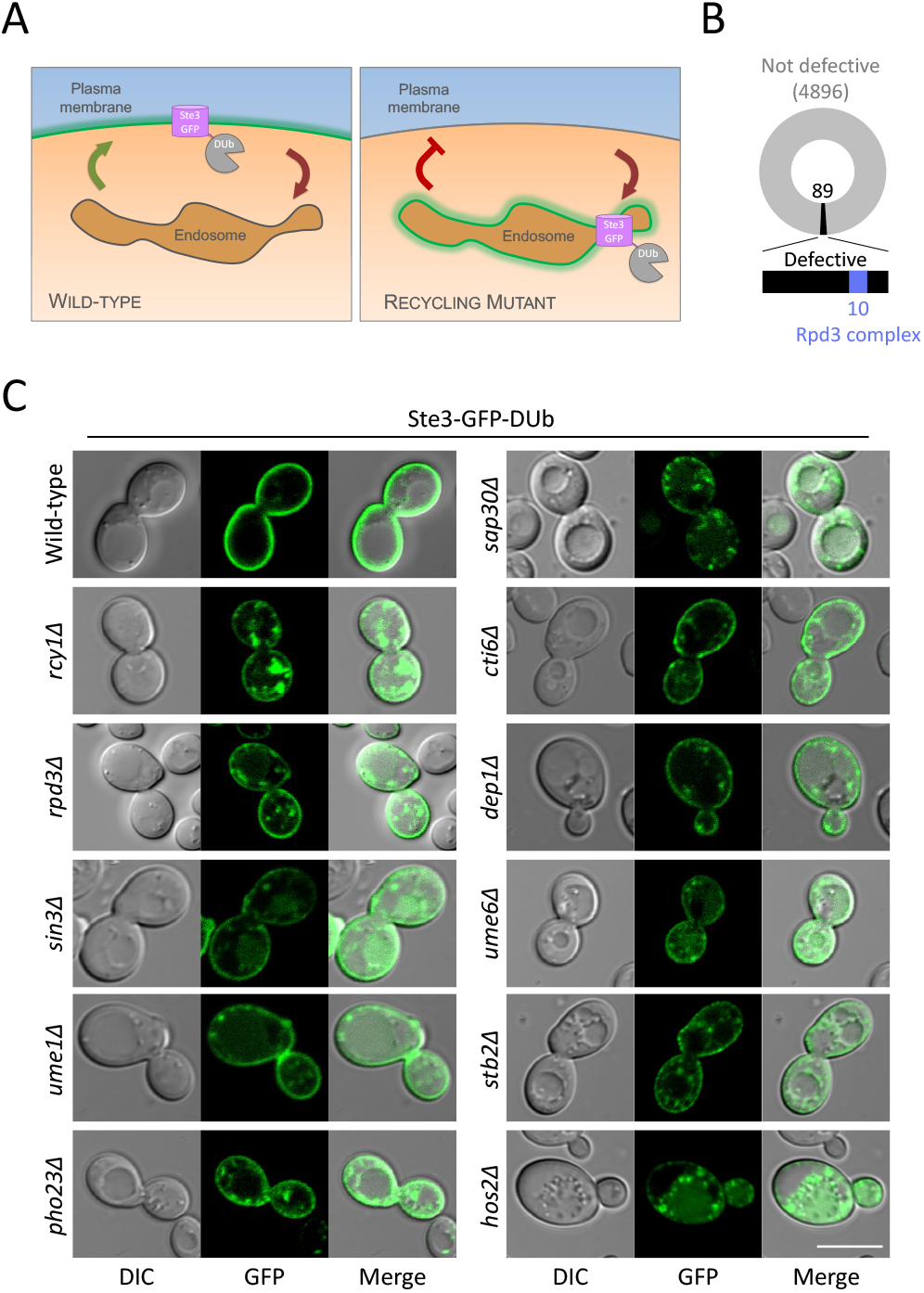
The Rpd3-complex is required for Ste3-GFP-DUb recycling. **A)** Schematic diagram showing Ste3-GFP-DUb recycling reporter, which efficiently recycles in wild-type cells (left) but accumulates in intracellular endosomal compartments in mutants with defective recycling (right). **B)** Results from localisation screen revealed 4,896 gene deletion mutant strains had no defect in recycling Ste3-GFP-DUb to the surface (grey) but 89 mutants were defective in recycling (black), including 10 members of the Rpd3 complex (blue). **C)** Wild-type and indicated mutant cells expressing a chromosomally integrated version of Ste3-GFP-DUb under control of the *STE3* promoter were grown to mid-log phase and imaged by confocal fluorescence microscopy. Scale bar, 5 μm.

**Figure 2:**
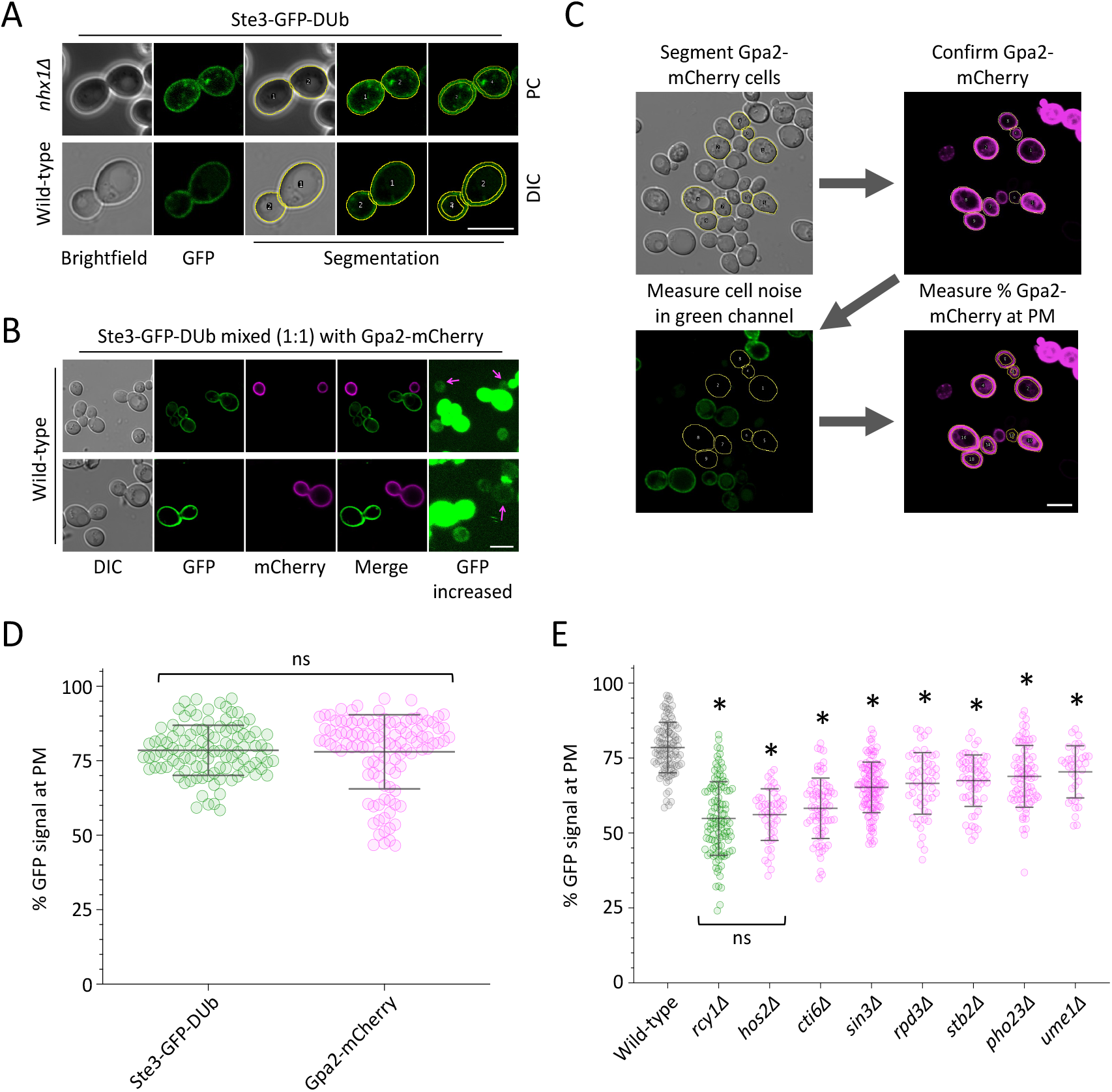
Quantification of Ste3-GFP-DUb recycling defects in Rpd3-complex mutants. **A)** Cells expressing Ste3-GFP-DUb were grown to mid-log phase and imaged with Phase Contrast (top) or DIC (bottom) objectives. Segmentation with ROI shown overlaid on each fluorescence image. A second ROI that excludes plasma membrane signal was created by morphological erosion. **B-C)** Wild-type cells expressing Gpa2-mCherry from the *CUP1* promoter, induced by addition of 100 μM copper chloride to the media, and wild-type cells expressing Ste3-GFP-DUb were mixed at a 1:1 ratio and imaged. mCherry expressing cells were used to identify and segment wild-type cells lacking GFP signal, allowing cellular background fluorescence in the green channel, observed at increased intensity (right), to be measured. **D)** The surface levels of Ste3-GFP-DUb and Gpa2-mCherry were both calculated as a percentage. **E)** The percentage of plasma membrane Ste3-GFP-DUb signal in wild-type cells (grey, n=104) was compared with *rcy1*Δ mutants (green, n =111) and various mutants (magenta) of the Rpd3 complex: *hos2Δ* (n=47), *cti6Δ* (n=73), *sin3Δ* (n=114), *rpd3Δ* (n=57), *stb2Δ* (n=61), *pho23Δ* (n=77), and *ume1Δ* (n=36). Student’s *t*-test comparisons between wild-type and each mutant were performed and asterisks (*) used to indicate significant difference (p < 0.0001). Scale bar, 5 μm.

To validate the role of the Rpd3 complex in surface recycling, we have previously employed an assay that takes advantage of recycling mutants exhibiting defective surface localisation of the tryptophan permease Tat2 (**Figure 3A**), which is required for growth in limited tryptophan media [47]. This assay requires tryptophan auxotroph strains, so we tested Rpd3-complex mutants in the SEY6210 background that harbours a *trp1-Δ901* mutation for growth capacity in media of replete (40 mg/L) versus low tryptophan concentrations (5 mg/L and 2.5 mg/L). We have recently documented a quantitative analysis method for such growth assays across large spectrum of serial dilutions [48], which we used to quantify growth defects attributed to defective Tat2 recycling in Rpd3-complex mutants (**Figure 3B, 3C**). We confirmed that most mutants had a low-tryptophan dependent phenotype. Importantly, only *pho23Δ* cells exhibited wild-type like growth in both low (5 mg/L and 2.5 mg/L) tryptophan media, and *ume1Δ* cells in 2.5 mg/L tryptophan had growth that was not significantly different to wild-type. Convincingly, *pho23Δ* and *ume1Δ* mutants were the two least defective mutants from our quantitative analysis of Ste3-GFP-DUb recycling in a completely different strain background. This indirect assay further suggests that Tat2 recycling is perturbed in all these mutants lacking Rpd3-subunits. We note that expression of *TAT2* in *rpd3Δ* cells is very similar to wild-type cells [49,50], but even when Tat2-3xHA was expressed from an endogenous promoter levels were significantly reduced in *rpd3Δ* cells (**Figure 3D, 3E**). We assume the reduced recycling of Tat2 that impairs tryptophan uptake results in increased trafficking of Tat2 from endosomes to the vacuole for degradation.

**Figure 3:**
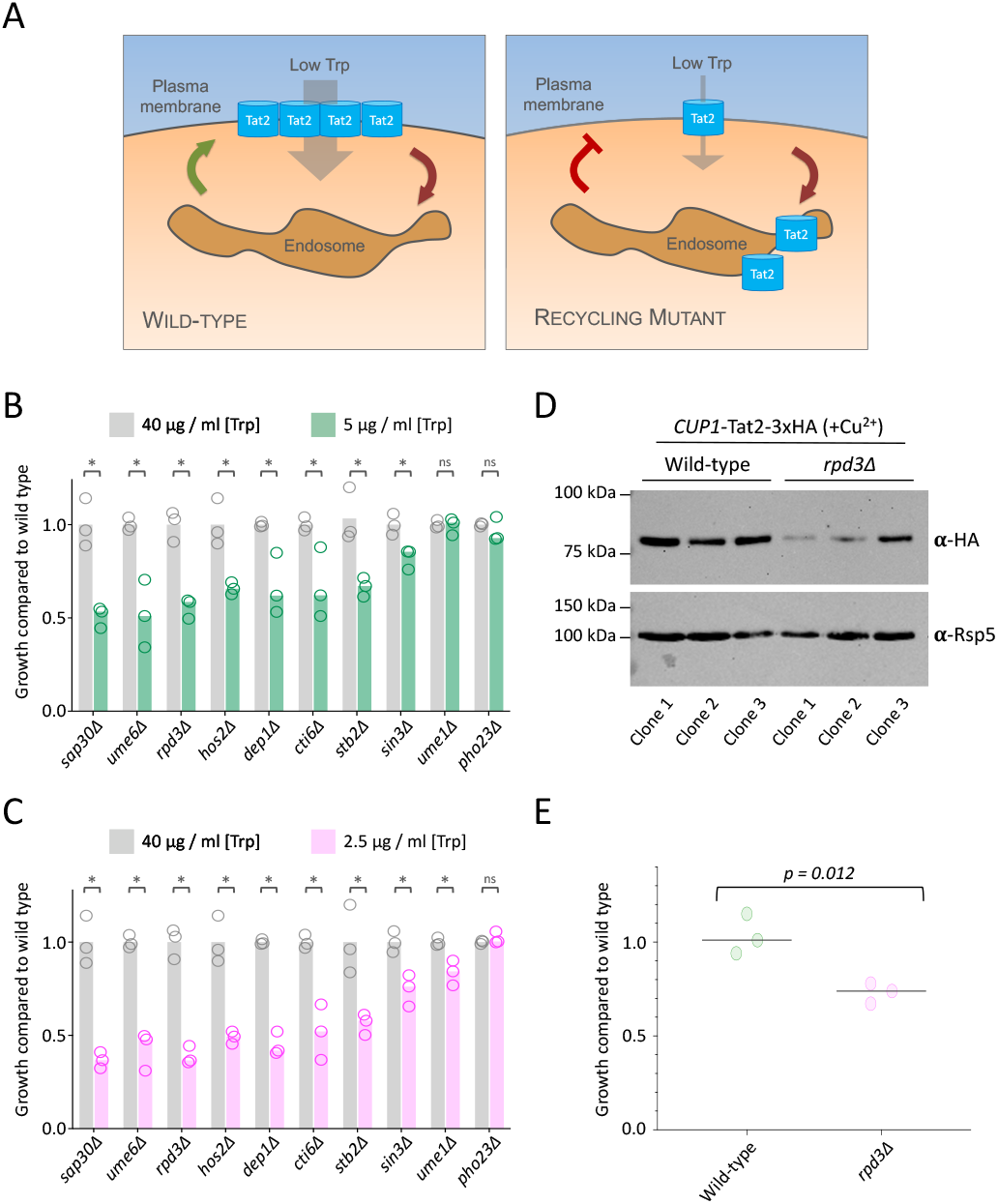
The Rpd3-complex is required for Tat2 recycling. **A)** Schematic diagram showing the uptake of tryptophan via the high affinity Tat2 permease. In tryptophan auxotroph cells grown on media containing low tryptophan concentrations, Tat2 uptake is required for efficient growth of wild-type cells (left) and Tat2 recycling defects inhibit growth (left). **B-C)** Yeast were grown to mid-log phase and spotted out on media of replete (40 μg/ml) and limited, either 5 μg/ml (in B) and 2.5 μg/ml (in **C)** tryptophan concentration. Growth was measured across multiple serial dilutions and calculated as a ratio compared to wild-type cells from the same plate. Asterisks (*) used to indicate significant difference (p < 0.03) from *t*-test comparisons. **D)** Cells transformed with a Tat2-3xHA plasmid containing a copper-inducible *CUP1* promoter were grown in media containing 50 μM copper chloride to mid-log phase before lysates were generated for immunoblot analysis using **α**-HA and **α**-Rsp5 antibodies (upper). **E)** Densitometry was used to measure the signal intensity of Tat2-HA in different clones and strains from **(D)**, normalized to loading control.

### Hypothesis for Rpd3-complex in recycling

Having confirmed and quantified that the requirement of the Rpd3 complex in surface recycling of diverse cargoes, we set out to confirm the mechanisms regulating this trafficking pathway. As previously documented, Rpd3 primarily localises to the nucleus [34] As expected, Rpd3 with a C-terminal mCherry tag (Rpd3-mCherry) localised to the nucleus labelled with Hoechst stain, but also to the cytoplasm (**Figure 4A**). Similar results were observed for an N-terminally tagged fusion of mCherry-Rpd3 (**Figure 4B**). The Ste3-GFP-DUb recycling defect of *rpd3Δ* cells was successfully complemented by expression of tagged Rpd3, showing these fusion proteins are functional (**Figure 4C**). The nuclear localisation of the KDAC Rpd3 allows regulation of transcription via the posttranslational modification of chromatin [31]. Rpd3 forms both small and large complexes with Sin3 and other subunits that contribute to its regulation [44,51], with various additional studies further documenting protein-protein interactions between Rpd3-complex subunits [52–56], which we highlight in a physical interaction network (**Figure 4D**). Importantly, many of these factors (10 out of 14) were independently identified from a blind genetic screen for recycling machinery [19] and subsequently validated and quantified (**Figures 1 - 3**). We hypothesised that the Rpd3-complex regulates expression of either specific recycling factor(s) identified from the Ste3-GFP-DUb localisation screen or regulates the expression of an unknown essential gene not represented in the library of viable haploid deletions used for this screen.

**Figure 4:**
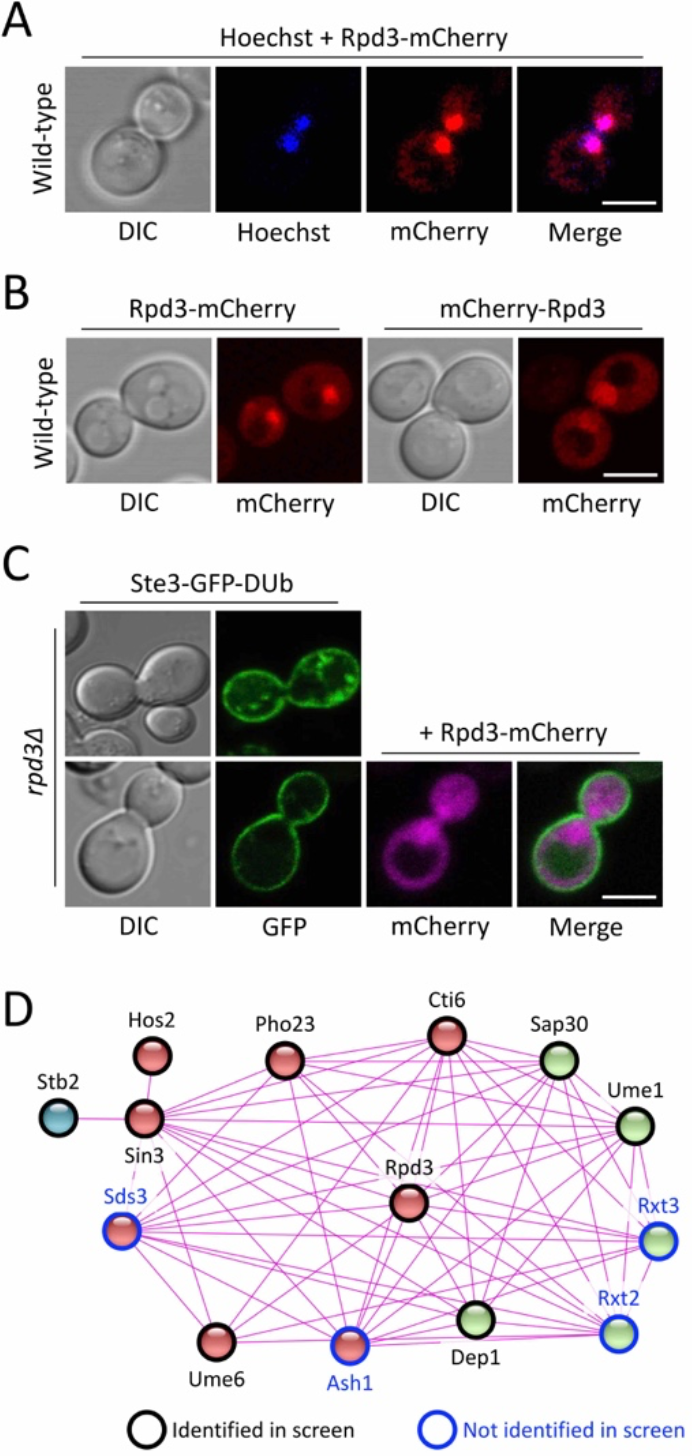
Rpd3 complex members regulate Ste3-GFP-DUb recycling. **A)** Yeast cells transformed with plasmids expressing Rpd3 from the *CUP1* promoter with either a C-terminal (left) or N-terminal (right) mCherry tag were grown to mid-log phase in media containing 50 μM copper chloride prior to confocal microscopy. **B)** Wild-type cells expressing Rpd3-mCherry were grown to mid-log phase, washed twice in fresh media and then incubated for 30 minutes with water containing 8 μM Hoechst, followed by fluorescence microscopy. **C)** Mutant *rpd3Δ* cells expressing an endogenously expressed version of Ste3-GFP-DUb were transformed with a vector control (upper) and a plasmid encoding Rpd3-mCherry under control of the *CUP1* promoter (lower). Transformants were grown to mid-log phase prior to confocal microscopy. **D)** A protein association network based only on experimental evidence of physical interactions (confidence = 0.400) was generated for the Large Rpd3 complex using STRING v11.5. Entries are coloured based on a kmeans clustering algorithm for 3 clusters (red, green, blue) but also outlines indicate whether mutants of these proteins were identified (black) or not (blue) from the Ste3-GFP-DUb localisation screen. Scale bar, 5 μm.

### Downstream Rpd3 targets regulate recycling

As recycling defects were phenocopied across most mutants lacking subunits of the Rpd3-complex, we reasoned that gene expression differences of potential target genes would be shared across mutants. Therefore, we assembled gene expression profiles [50] for the 89 validated recycling factors [19], depicted as a heatmap (**Figure 5A**) but averaged the changes in gene expression across all mutant conditions. This allowed us to identify 24 genes with significantly reduced expression across mutants. These represented our initial candidates for a complementation screen, as we hypothesised their repression via Rpd3-complex results in recycling defects. Therefore, reintroducing high levels of these factors might supress these recycling defects. We also included the next 10 genes that were decreased to a smaller degree, to account for gene regulation that was repressed significantly in only certain mutants (for example to account for technical errors during transcriptomic analyses). Each of these genes were over-expressed from a plasmid library [57] in *rpd3Δ* cells stably expressing Ste3-GFP-DUb. We were unable to test complementation of *GPA1* and *HDA1* as these clones repeatedly did not yield any transformations, potentially as the combinations of these over-expressors with *rpd3Δ* are not viable. We assessed reporter localisation in each over-expression condition from 3 independent transformants and used a qualitative scoring system to document results (**Figure 5B**). This screen revealed 3 factors, Sit4, Dit1 and Ldb7, that complement the recycling defect of *rpd3Δ* cells, which were all confirmed by further imaging experiments (**Figure 5C**) and quantification using the analysis pipeline discussed above (**Figure 5D**). We included images and analysis of Prm8 as a control, as *PRM8* was the most repressed gene that failed to complement recycling upon over-expression. To validate these complementation factors, we employed a distinct recycling assay based on the lipid dye FM4-64, which can be loaded to endosomes for brief labelling periods, followed by tracking efflux via recycling that triggers dye quenching [20]. We found that the rate of efflux from recycling mutants is reduced (**Figure 6A**). Efflux can be tracked through a kinetic assay using flow cytometry, measuring fluorescence from ~1500 cells per second and the fluorescence of each cell is presented as a percentage of the initial fluorescence, calculated as the average from all events in first 10 seconds (**Figure 6B**). This assay was used to show that plasmid over-expression of Sit4, Dit1 and Ldb7 all increase the rate of FM4-64 recycling observed in *rpd3Δ* cells (**Figure 6C-F**). A wild-type positive control, and *rpd3Δ* cells transformed with an empty vector as a negative control were performed at the same time and the profiles of these efflux measurements overlaid to each complementation profile.

**Figure 5:**
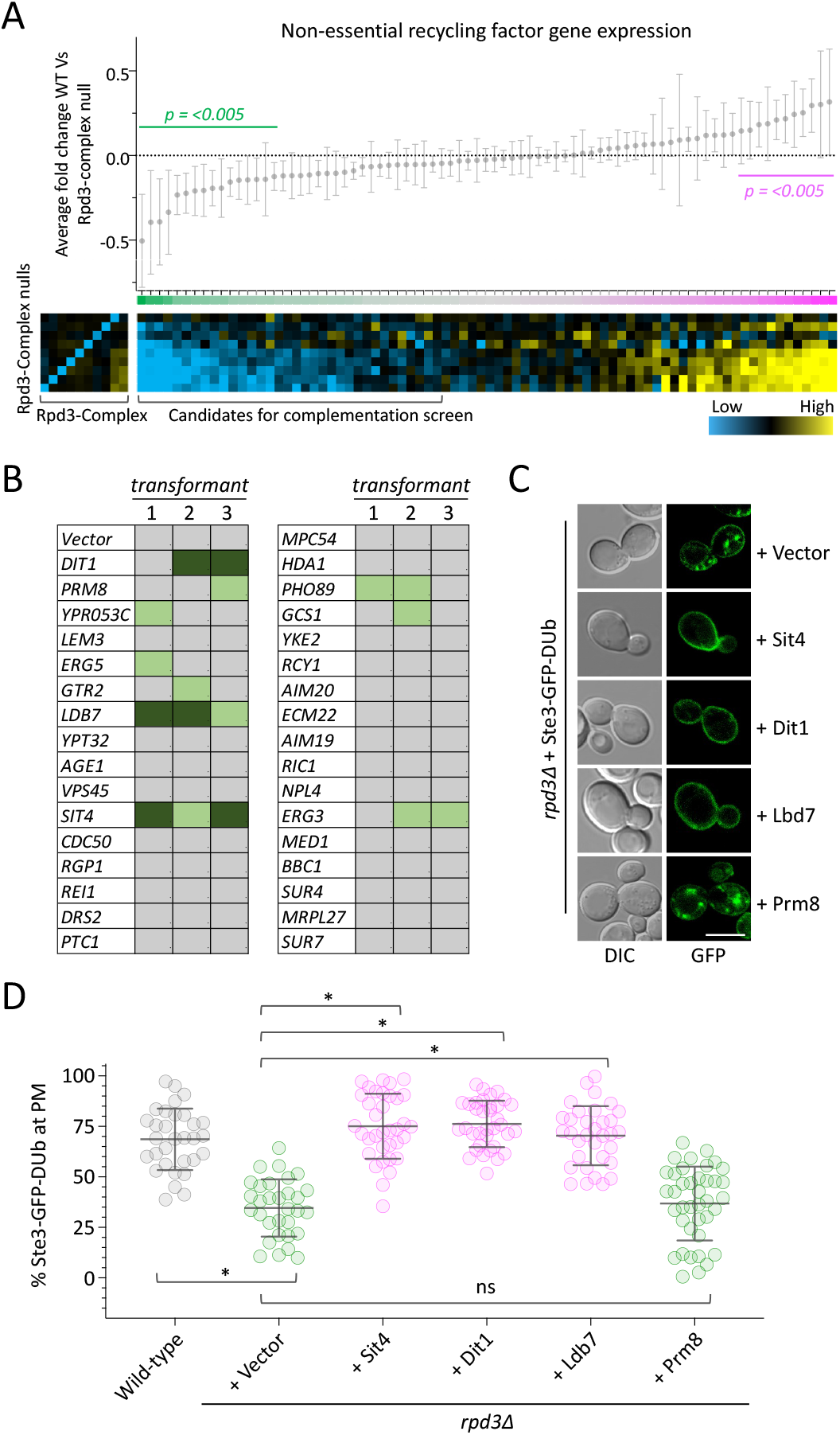
Complementation screen reveals Rpd3 recycling targets. **A)** Changes in gene expression of 89 validated recycling factors were averaged across various null mutants of the Rpd3-complex compared with wild-type, with mean ± standard deviation plotted (upper). Individual log2 fold-change expression profiles were also assembled (organised rows top - bottom: *stb2Δ*, *ume1Δ*, *hos2Δ*, *cti6Δ*, *pho23Δ*, *sap30Δ*, *dep1Δ*, *sin3Δ*, *rpd3Δ*) as a heat map (lower right). The successful deletion of each gene is shown to reduce expression of each individual subunit (lower left). **B)** Each of the listed genes were chosen for over-expression in *rpd3Δ* cells stably integrated with Ste3-GFP-DUb. Single colony transformants of each were grown to mid-log phase and imaged by confocal microscopy. Transformants that rescue Ste3-GFP-DUb recycling were first scored qualitatively, with modest (light green) and substantial (dark green) levels of potential complementation of recycling indicated. **C)** Over-expression candidates identified from the screen described in **(B)** were grown to mid-log phase and prepared for confocal imaging. **D)** Quantification of surface level Ste3-GFP-DUb in each of the indicated cellular conditions, with asterisks (*) used to indicate significant difference of p < 0.0001 from *t*-test comparisons. Scale bar, 5 μm.

**Figure 6:**
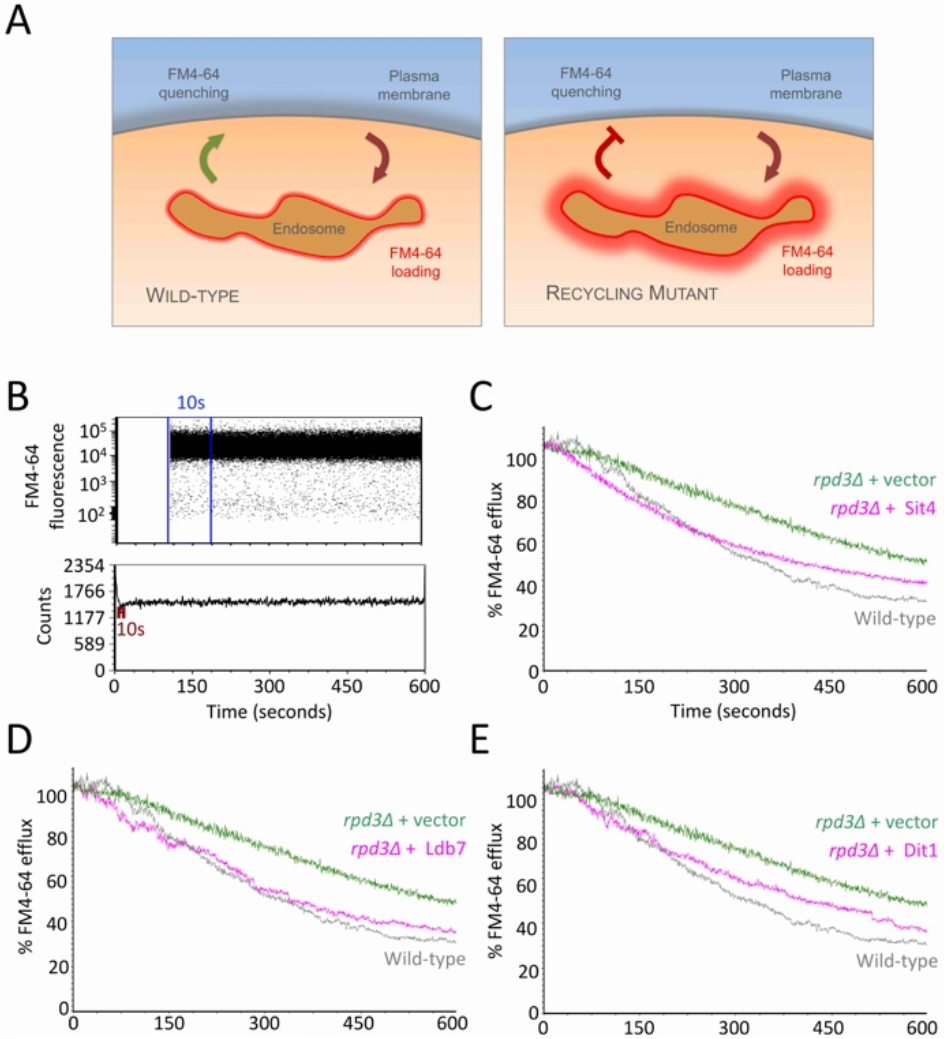
Over-expression of Sit4, Ldb7 and Dit1 rescues FM4-64 recycling defect of *rpd3Δ* mutants. **A)** Schematic representation of dye recycling assay, where the fluorescent lipid dye FM4-64 is loaded to endosomes for 8 minutes at room temperature in YPD media containing 40 μM FM4-64, subjected to 3x 3-5 minute washes in ice cold minimal media prior to a small volume of washed culture (~5 - 15 μl) brought up in 3 mls media maintained at room temperature followed by flow cytometry measurements. **B)** FM4-64 fluorescence is measured by flow cytometry and the average fluorescence measured across the first 10 seconds and use to calculate all further measurements as a percentage of this average (upper). Voltage and flow rate are set to analyse 1000 - 3000 cells per second, with measurements acquired for 10 minutes total (lower). **C - E)** Efflux measurements following protocol in **(A)** were acquired for *rpd3Δ* cells transformed with plasmids over-expressing Sit1 **(C)** Ldb7 **(D)** and Dit1 **(E)**. As a control, the efflux profile of wild-type and *rpd3Δ* cells labelled and measured during the same session are included in each graph.

### Rpd3 regulates the essential factor Pik1

As the Rpd3 complex might also regulate expression of essential genes that are involved in membrane trafficking of surface proteins. To explore this possibility, we used recently optimised bioinformatics approaches [48] to assemble gene expression profiles of only essential genes that were not tested in the original recycling reporter screen (**Figure 7A**). We compared essential gene profiles of 7 different null strains representing Rpd3-complex members [50], many of which share large regions of expression patterns with *hos2Δ* and *ume1Δ* cells being most distinct (**Figure 7B**). As the recycling phenotypes are shared across the various mutants, we averaged the changes in expression across mutants to identify those with most significantly altered expression. Gene ontology analyses of the most repressed 43 genes (log2 fold change < 5.0) was performed and showed enrichment for processes including autophagy, phosphorylation and lipid regulation (**Figure 7C**), with a large amount of annotation overlap of the most enriched (**Figure 7D**). We considered *PIK1*, a phosphatidylinositol-4-kinase (PI4K) that regulates trafficking from both the Golgi and endosomes [58], would be a likely Rpd3 target gene with potential to regulate trafficking of surface membrane proteins. Indeed, the levels of *PIK1* are substantially decreased in mutants of the Rpd3-complex, even when viewed with all essential and non-essential gene profiles (**Figure 7E**). To test the hypothesis that Pik1 is required for efficient recycling, we again employed the FM4-64 efflux assay (**Figure 6A**). For this, wild-type cells and temperature sensitive mutants were loaded with FM4-64 and the rate of efflux was measured over time. There was a significant decrease in recycling efficiency in cells expressing either mutant allele of *pik1* (*pik1-83* and *pik1-139*), even when the cytometry experiments performed in media that was not at restrictive temperature (**Figure 7F**).

**Figure 7:**
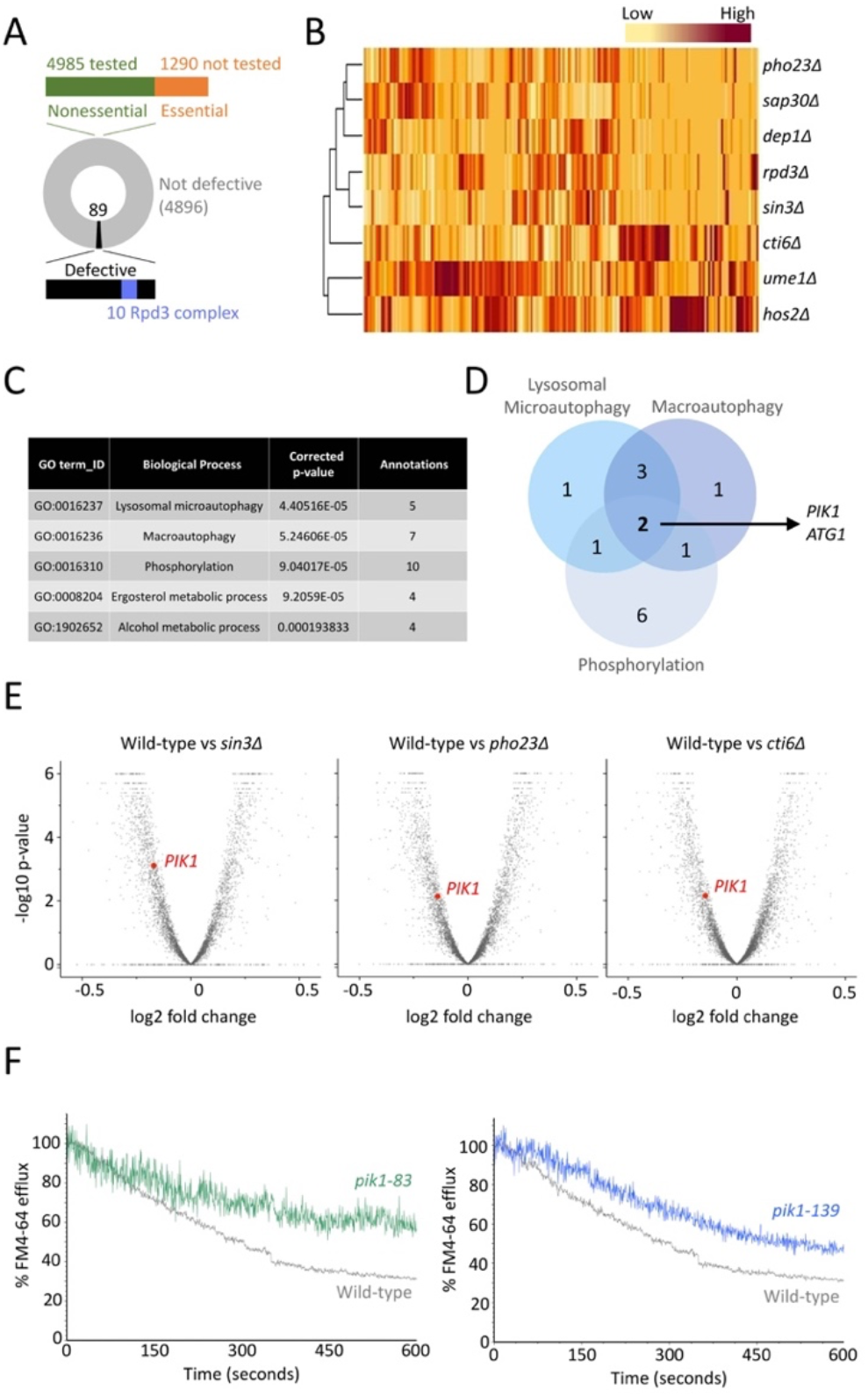
The essential *PIK1* gene is an Rpd3 target that regulates surface recycling. **A)** Pictorial representation of Ste3-GFP-DUb localisation screen where 4,985 nonessential mutants were tested (green) but the 1,290 essential genes were not (orange). **B)** Heat map generated from changes in essential gene expression when indicated mutants are compared with wild-type cells. **C)** Gene Ontology analysis was performed on essential genes that had reduced expression of 5-fold or more when different *rpd3Δ* mutants **(B)** on average across all mutants (totalling 43 genes). **D)** Venn diagram showing the overlap of GO annotations **(C)** in top three scoring biological processes. **E)** Volcano plot of all genes, including essential and non-essential ORFs, showing changes in gene expression compared with wild-type cells for *sin3Δ*, *pho23Δ*, and *cti6Δ*. The reduced levels of *PIK1* are highlighted in each comparison (red). **F)** FM4-64 efflux measurements of wild-type cells and mutants harbouring temperature sensitive alleles of *pik1 p*(*ik1-83*, green and *pik1-139*, blue) were performed by flow cytometry and plotted as a % of average initial fluorescence (from first 10 seconds).

It has previously been proposed that the unfolded protein response (UPR), which is elevated in *rpd3Δ* cells, results in surface proteins like the uracil permease Fur4 to be retained in the ER and degraded [41]. This conclusion was based on dramatically reduced cellular levels of fluorescently tagged Fur4, not increased levels of GFP retained following vacuolar degradation. However, Fur4 tagged with mNeonGreen (mNG) expressed in wild-type and *rpd3Δ* cells localises at the PM and inside the vacuole, with no indication of ER retention (**Figure 8A**). Furthermore, we observed no difference in overall levels of Fur4-mNG between wild-type cells and *rpd3Δ* mutants (**Figure 8B**). For this reason, we propose the regulation of Rpd3 on cell surface proteins is not indirect via the UPR, but through regulation of factors required for recycling. We conclude that both non-essential and essential gene targets of the Rpd3-complex have the capacity to regulate cell surface recycling governed at the transcriptional level (**Figure 8C**).

**Figure 8:**
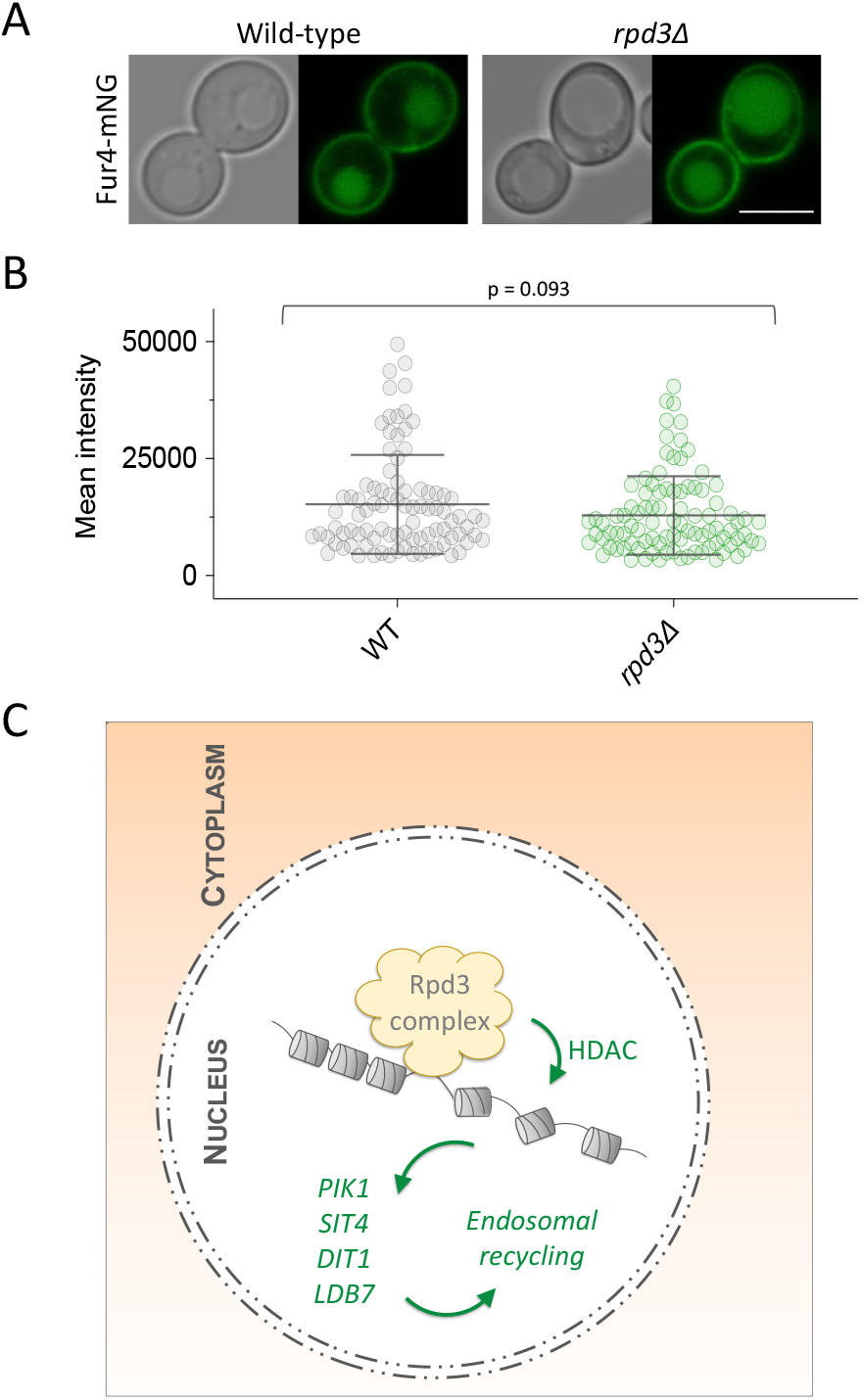
Summary model. **A)** Wild-type and *rpd3Δ* cells expressing Fur4-mNeonGreen (Fur4-mNG) from a plasmid were grown to mid-log phase, resuspended in azide containing buffer and then imaged using confocal fluorescence microscopy. **B)** Micrographs from **(A)** were segmented based on DIC images and then the GFP fluorescence signal measured for each wild-type (n = 93) and *rpd3Δ* (n = 94) cells, then mean intensity plotted. **C)** The Rpd3 chromatin remodelling complex localises to the nucleus and post-translationally deacetylates histones (HDAC activity) to control expression of *PIK1*, *SIT4*, *LDB7* and *DIT1*, which are all required for efficient cell surface recycling of internalised surface membrane proteins and lipids. Scale bar, 5 μm.

## DISCUSSION

The lysine deacetylase Rpd3, alongside many of its physical interactors, are known to have massive effects on gene expression in yeast [50,59–61]. The identification of all 10 members of the complex from a blind screen [19] strongly implicate the Rpd3 complex as a regulator of recycling in yeast. In this study, we showed that these 10 subunits are required for recycling, but to different degrees. We stably integrated Ste3-GFP-DUb to give consistent phenotypes over plasmid-borne expression initially used to screen for factors. This allowed a quantification approach to specifically measure how much GFP signal was found efficiently recycled to the PM versus signal retained in endosomes (**Figure 2**). Importantly, these results were compared to the quantitation of a growth defect indirectly associated with Tat2 recycling (**Figure 3**). For example, *hos2Δ* mutants were amongst the most defective mutants in both quantified assays, and both *ume1Δ* and *pho23Δ* were the least defective. We note that unlike Ste3-GFP-DUb localisation, the tryptophan-uptake assay was not sensitive enough to identify defects in Tat2 recycling in *pho23Δ* mutants. Although Rpd3 is associated with global deacetylation events [31], Pho23 is more limited to a specific subset of loci [62,63], so it may be that Pho23 does not regulate all genes associated with efficient recycling. We show the DNA-binding protein Ume6 is required for recycling, with *ume6Δ* mutants one of the most defective Tat2 recycling mutants. The genetic screen did not identify several Rpd3 subunits, such as the DNA-binding protein Ash1 (**Figure 4**), which exhibits gene specific regulation with Ume6 [64]. Therefore many, but not all, Rpd3-subunits are required for efficient transcriptional control of the recycling pathway.

To explain these results, a phenotypic complementation screen was performed by over-expressing downstream recycling genes repressed in *rpd3Δ* mutants. We predicted this over-expression strategy using 2μ based plasmids retained with 100s of copies per cell [65], would overwhelm repression mechanisms in *rpd3Δ* cells to reveal and *bona fide* target recycling genes. 32 candidates were tested in this screen, with only two additional candidates, *HDA1* and *GPA1*, failing to yield viable transformants after various attempts and optimisations. Higher levels of Hda1, which is histone deacetylase related to Rpd3 with shared molecular activity [31], might induce lethality in *rpd3Δ* mutants. This screen revealed 3 validated hits that rescue recycling of *rpd3Δ* mutants: the protein phosphatase Sit4, the transcriptional regulator Ldb7 and the sporulation factor Dit1 (**Figures 5 & 6**). The Sit4 phosphatase is a strong candidate for regulating recycling as it has been previously shown to modify machinery in secretory [66] and endocytic [67] trafficking pathways. Ldb7 is itself a transcriptional regulator [68], in the family of low-dye-binding mutants associated with Golgi function, stress response and cell wall organisation [69], any of which might indirectly impinge in recycling. Finally, Dit1 which regulates formation of spore walls following developmental expression of starved diploid cells [70,71]. *DIT1* is repressed in haploids via Ssn6-Tup1 repressor [72] and increased greatly during sporulation [73]. However, many independent studies report haploid expression of *DIT1*, that can be further repressed upon increased temperature [74], deletion of *RPD3* [49] or addition of the Rpd3 inhibitor trichostatin A [75], potentially pointing to a distinct function. Finally, the implication of the Pik1 in recycling (**Figures 7**) is easily rationalised as it is known to modify lipids required for proper Golgi and endosomal trafficking [58], so fine tuning of these various pathways that control surface proteins could be mediated via the Rpd3-complex.

Our observations are consistent with other reports in the literature relating to surface protein regulation and the Rpd3-complex. For example, the Trk2 surface potassium channel has been proposed to be negative regulated in cells mutants of *rpd3* [76]. Similarly, although as *rpd3Δ* cells have increased expression of the major acid phosphatase *PHO5* [77], phosphate uptake via the surface localised H^+^/PO_4_^3−^ symporter Pho84 is defective in *rpd3Δ* cells, with accumulation of Pho84-GFP in endosome-like compartments in cells lacking *RPD3* [78]. Encouragingly, both our bioinformatic and functional observations on recycling align with a study on how yeast cells respond to exposure to the anti-malaria drug artemisinin, which found both *rpd3Δ* and *sit4Δ* cells were hypersensitive to the drug and suggest this is due to premature degradation of surface proteins [41]. Beyond this, we have previously shown that the developmentally regulated expression of the Cos proteins, which drive ubiquitin-mediated vacuolar degradation of surface proteins *in trans* [79], is ablated by deletion of either *RPD3* or *SIN3* [80]. This demonstrates that complex and overlapping modes of transcriptional regulation control the membrane trafficking routes used by cell surface membrane proteins.

The experimental validation of these candidates demonstrates the complexity of surface protein recycling in yeast. As Rpd3 and some of the downstream recycling gene targets discussed are highly conserved throughout evolution, this regulatory control could be maintained in other eukaryotic systems. Indeed, Rpd3 has orthologues expressed in various other eukaryotic systems, and its roles could be understood in terms of regulating surface membrane proteins. For example, in Drosophila, Rpd3 is required for cells to respond appropriate to nutrient starvation [81]. For discoveries in yeast relating to recycling and its metabolic or transcriptional control, we recommend using more than one assay, such as Ste3-GFP-DUb localisation (**Figure 1**), Tat2 mediated tryptophan update (**Figure 3**), or efflux of internalised FM4-64 (**Figure 6**), to fully validate any defect. Beyond this, we promote the use of bioinformatics to identify downstream targets of transcriptional regulators for experimental testing, as discussed in this study. Future work will be aimed at deciphering the individual roles of these new candidates in the recycling pathways and understanding any functional overlap.

## METHODS

### Reagents

Yeast strains and are included as supplemental tables (**Table S1** and **Table S2**, respectively).

### Yeast strains and culture conditions

Yeast strains used in this study are listed in **Table S1** and were grown in either yeast extract peptone dextrose (YPD) media, for example when making competent cells, or synthetic complete (SC) minimal drop-out media lacking appropriate bases/amino acids, when selection of plasmids or integrations was necessary. Competent yeast stocks were prepared in Li-TE sorbitol buffer (100mM lithium acetate, 10mM Tris.HCl pH 7.5, 1.2M sorbitol, 1mM EDTA, 200μM calcium chloride) and plasmids incubated for 40 mins at 30°C followed by heat shock at 42°C for 20 minutes and plating on solid selective media. Yeast cultures were prepared by inoculation from a clonal yeast patch and grown overnight at 30°C in 5 ml two-fold serial dilutions to ensure cells used in experimental procedures were at mid log (OD_600_ = ~1.0). Expression of proteins from the *CUP1* promoter was induced by addition of 50 μM Copper Chloride (CuCl_2_) to the media for at least 1 hour prior to experiments. Nuclei of yeast cells were labelled with fluorescent DNA stain by first growing to mid-log phase, washing in fresh SC media, prior to addition of 8 μM Hoechst-33342 (Invitrogen−) for 30 minutes.

### Bacterial culture

Plasmid DNA listed in **Table S2** was stored and propagated in Top10 *Escherichia coli* (Invitrogen−). For plasmid isolation, *E. coli* were grown in 2YT media (w/v: 1.6% tryptone, 1% yeast extract, 0.5% NaCl) containing either 100 μg/ml ampicillin sodium salt (Melford) or 50 μg/ml kanamycin monosulphate (Formedium) each diluted from 1000x frozen stock.

### DNA manipulations

Yeast expression plasmids were purified from ~5ml saturated cultures using a Wizard^®^ DNA Purification System (Promega) and were transformed into competent yeast. The Gibson assembly principle [82] of incubating homologous PCR products with Taq ligase (NEB), T5 exonuclease (NEB) and Phusion polymerase (NEB) for 1 hour at 50°C, followed by plating on selective 2YT media, was used to create different fluorescently labelled Rpd3 expression constructs, which were confirmed by Sanger sequencing. Stable integrations of Ste3-GFP-DUb under control of the *STE3* promoter were performed by linearising pCM850 with NsiI followed by ethanol precipitation and transformation into the various parental yeast strains. For strains with *loxP* flanked integrations, cassettes were excised using a modified *TEF1-Cre* expression system [83]

### Fluorescence microscopy

Yeast cells were grown to mid-log phase and prepared for confocal microscopy by centrifugation to concentrate samples and resuspension in water prior to storage on ice prior to confocal microscopy. Imaging was performed from live cells using a laser scanning confocal microscope (Zeiss LSM 780) with a 63x Differential Interference Contrast (DIC) or 63x Phase-Contrast (PC) oil-immersion objectives (Objective Plan-Apochromat 63x/1.4 Oil, Numerical Aperture 1.4, Zeiss). Fluorescence microscopy images were captured via Zen Black (Zeiss) software and modified for contrast, colour and merge in ImageJ (version 2.0.0).

### Flow cytometry FM4-64 recycling assay

Mid-log phase yeast cells were concentrated 5-10x and brough up in fresh 100 μl YPD containing 40 μM FM4-64 dye (N-(3-Triethylammoniumpropyl)-4-(6-(4-(Diethylamino) Phenyl) Hexatrienyl) Pyridinium Dibromide) dye. FM4-64 was loaded to endosomes for 8 minutes at room temperature prior to 4x 5minute washes in cold SC media. Cells were resuspended in a small volume (100 - 200 μl) ice cold SC media and ~10 μl added to a flow cytometer tube with 3 mls room temperature SC media and fluorescence measurements taken immediately for a 600 second period using an LSR Fortessa instrument (BD Biosciences). Flow rate was set to flow at a rate between 1000 – 2000 cells per second. FM4-64 fluorescence was measured with a 561nm excitation laser, and emission filter 710 / 50. Flow data was analysed using FCS Express (version 7.06.0015; DeNovo).

### Immunoblotting

Lysates were generated from mid-log phase yeast by resuspension in 0.2 M sodium hydroxide for 3 minutes before pelleting and resuspension in TWIRL buffer (8M urea, 10% glycerol, 5% SDS, 10% 2-Mercaptoethanol, 50mM Tris.HCl pH 6.8, 0.1% bromophenol blue). Lysates were resolved by SDS-PAGE before protein transferred to nitrocellulose using the iBlot2 system (ThermoFisher). Membrane was blocked in 5% milk followed by probing with either α-HA Mouse Monoclonal (Catalogue #HA.11; Biolegend, SanDiego, CA) or α-Rsp5 Rabbit Polyclonal [84] antibodies. Secondary antibodies conjugated to HRP (Abcam, PLC) were used to visualise signals using the Pico Plus (ThermoFisher) Enhanced chemiluminescence substrate and an iBright™ Imager (ThermoFisher).

### Bioinformatics & statistics

For non-essential (**Figure 5**) and essential (**Figure 7**) recycling gene candidates, gene expression profiles were averaged across all mutants, since all mutants phenocopy one another with regards to recycling defects, and prioritised based on this average. Heat maps were generated to show distribution of individual values across mutants. Microarray data documenting expression changes (log fold and p values) were read into RStudio (version XXX RStudio Team, 2020) then processed using the dplyr (v1.0.7; Wickham et al., 2021) and tidyverse (v1.3.0; Wickham et al., 2019) packages to include only data for indicated deletion strains of the Rpd3-complex. The data were further sub-setted to include genes deemed essential for viability (1290 ORFs represented) acquired from the Saccharomyces Genome Database. Hierarchical clustering was visualised in the form of a heatmap using base R. Gene Ontology enrichments were performed using GO Term Finder (version 0.86) via YeastMine [85,86]. Physical interaction maps were generated using STRING pathway (version 11.5) analysis software [87]. Statistical analyses were performed using Graphpad (Prism, version 9.0.2).

## ACKNOWLEDGMENTS

Thanks to the York Bioscience Technology Facility imaging team for help with imaging and cytometry. This research was supported by a Sir Henry Dale Research Fellowship from the Wellcome Trust and the Royal Society 204636/Z/16/Z (CM). We are grateful to Paul Pryor (University of York) for access to a yeast 2μ plasmid over-expression collection. Thanks also to Chris Stefan (University College London) for the gift of yeast strains harbouring mutant alleles of *pik1*.

